# Bedaquiline Amplifies Proteasome Inhibitor Efficacy and Overcomes Resistance in Multiple Myeloma

**DOI:** 10.1101/2025.06.29.661768

**Authors:** Michela Cumerlato, Monica Maccagno, María Labrador, Beatrice Luciano, Elisabetta Mereu, Mariangela Porro, Cecilia Bandini, Domenico Maisano, Mattia D’Agostino, Alessandra Larocca, Francesca Gay, Annalaura Tamburrini, Francesca Anselmi, Paolo Ettore Porporato, Salvatore Oliviero, Benedetto Bruno, Eugenio Morelli, Nikhil Munshi, Roberto Piva

## Abstract

Proteasome inhibitors (PIs) are cornerstone therapies for multiple myeloma (MM), yet resistance remains a major barrier to durable responses. To identify druggable vulnerabilities that enhance PIs efficacy, we performed a small-molecule chemical screen in the presence of carfilzomib (CFZ). We identified bedaquiline (BDQ), an FDA-approved antimycobacterial agent, as a potent synergistic partner. BDQ and its fumarate salt (BDQ-F) significantly amplified CFZ-induced cytotoxicity in PI-sensitive and PI-resistant MM cells, in AL amyloidosis and other B-cell malignancies, with minimal toxicity toward normal cells. Mechanistic studies confirmed that BDQ specifically targets the ATP5F1C subunit of mitochondrial ATP synthase. BDQ-CFZ combination triggered extensive apoptosis, exacerbating proteotoxic stress and proteasome-associated pathways. BDQ specifically enhanced CFZ’s inhibition of the proteasome’s chymotrypsin-like activity. Importantly, BDQ synergized with multiple proteasome and ubiquitin-activating enzyme inhibitors, but not with other standard MM agents, underscoring its selective interaction with the UPS pathway. BDQ-CFZ co-treatment markedly reduced MM cell viability and tumor burden in patient-derived cells and zebrafish xenograft models. These findings support the therapeutic repurposing of BDQ to potentiate PIs efficacy and overcome resistance in MM and related B-cell malignancies.

**KEY POINTS:**

- Bedaquiline synergizes with proteasome inhibitors in MM and other hematologic malignancies, sparing normal cells
- Bedaquiline targets ATP synthase γ and boosts carfilzomib by enhancing chymotrypsin-like proteasome inhibition

**NOVELTY:** We identify the antimicrobial bedaquiline and its fumarate salt as potent enhancers of proteasome inhibitor efficacy in multiple myeloma and other B-cell malignancies. By inhibiting ATP synthase γ, bedaquiline amplifies chymotrypsin-like proteasome inhibition and overcomes drug resistance. This study reveals a mitochondria– proteasome vulnerability and proposes a clinically actionable strategy to restore sensitivity and reduce toxicity of proteasome-based therapies.

**VISUAL ABSTRACT:** 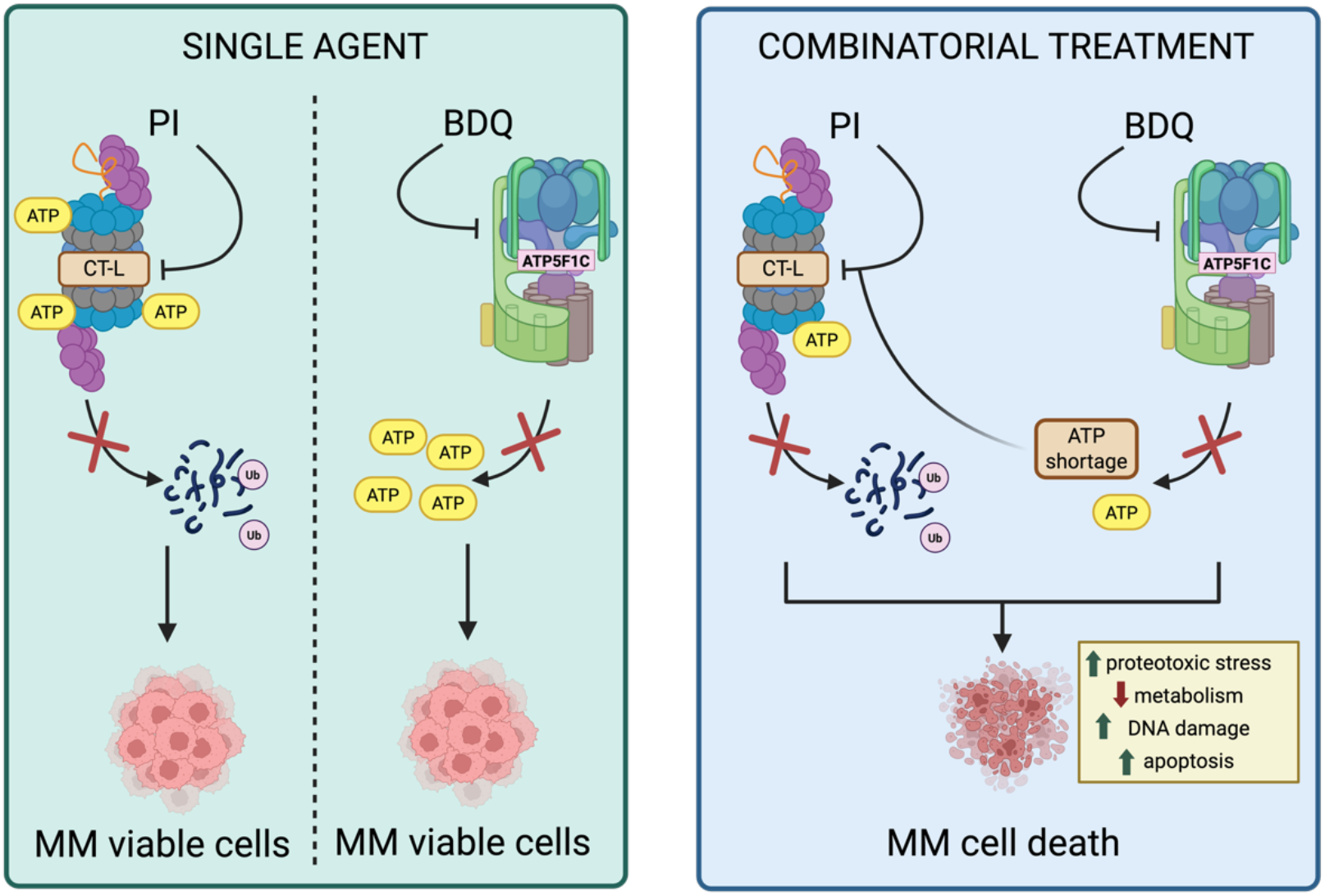

## INTRODUCTION

Multiple myeloma (MM) is a malignancy of terminally differentiated B cells marked by clonal plasma cell expansion in the bone marrow and excess monoclonal immunoglobulin production.^1^ It is the second most common hematologic cancer in high-income countries, typically diagnosed between ages 60 and 70.^2^ Standard frontline therapies combine proteasome inhibitors (PIs), immunomodulatory agents, corticosteroids, and monoclonal antibodies targeting CD38.^3–5^ Recently, CAR T cells and bispecific T cell engagers (BiTEs) have shown remarkable efficacy in heavily pretreated relapsed/refractory MM patients.^6,7^While these regimens have improved survival, MM remains incurable due to intrinsic and acquired resistance mechanisms.^8,9^Identifying new vulnerabilities and combinatorial strategies to enhance efficacy while reducing toxicity is essential.^10^ However, MM’s extensive genetic and epigenetic heterogeneity complicates treatment and fuels resistance.^11,12^To address this, omics-driven approaches and ex vivo functional screens are increasingly used to identify actionable targets.^13^ MM cells exhibit high proteostasis stress due to their immunoglobulin load, making them susceptible to PIs.^14^ Yet, metabolic and epigenetic reprogramming enables adaptation and drug resistance.^15,16^ Targeting these compensatory pathways offers a rational strategy to resensitize cells to PIs. We previously identified several PIs synergistic interactors, including IDH2 and LSD1, as actionable nodes to restore PIs sensitivity.^17–19^ Here, we describe the FDA-approved ATP synthase inhibitor bedaquiline (BDQ)^20^ as a novel synergistic partner of the PI carfilzomib (CFZ). BDQ-CFZ co-treatment selectively induces proteotoxic stress and apoptosis in MM cells. Given BDQ’s established safety and clinical availability, our findings support its repurposing to enhance PIs efficacy and overcome resistance in MM and potentially other B-cell malignancies.

## MATERIALS AND METHODS

Detailed experimental procedures for cell culture conditions, reagents, drug screening, cellular and biochemical assays, plasmids, virus production, RNA extraction, reverse transcription-quantitative PCR, western blotting, RNA sequencing, proteomics, data analysis, zebrafish xenograft assay, healthy donor and MM patient samples are included in Supplemental Material and Methods.

## RESULTS AND DISCUSSION

To identify compounds that enhance carfilzomib (CFZ) efficacy in multiple myeloma (MM), we performed a functional screen of 320 small-molecule inhibitors in the MM PI-resistant cells U-266^PIR^, in the presence or absence of sublethal CFZ dose (Fig. 1A). Top candidates were validated in five MM cell lines. Bedaquiline (BDQ) and its fumarate form (BDQ-F), approved to treat multidrug-resistant tuberculosis, showed strong synergy with CFZ in 5 and 3 out of 5 lines, respectively (Suppl. Table S1), and were prioritized for further study. BDQ and BDQ-F strongly synergized with CFZ in almost all tested PI-resistant and PI-sensitive MM cell lines (Fig. 1B–C; Suppl. Fig. 1A–B). The combination also enhanced cytotoxicity in systemic AL amyloidosis, mantle cell lymphoma, diffuse large B-cell lymphoma, and Burkitt’s lymphoma (Fig. 1D). Importantly, the BDQ-CFZ combination showed no synergistic toxicity in peripheral blood mononuclear cells (PBMCs) from healthy donors, immortalized B-lymphocytes (IST EBV TW6B), and bone marrow-derived stromal cells (HS-5) (Fig. 1E-G).

**Figure 1.**
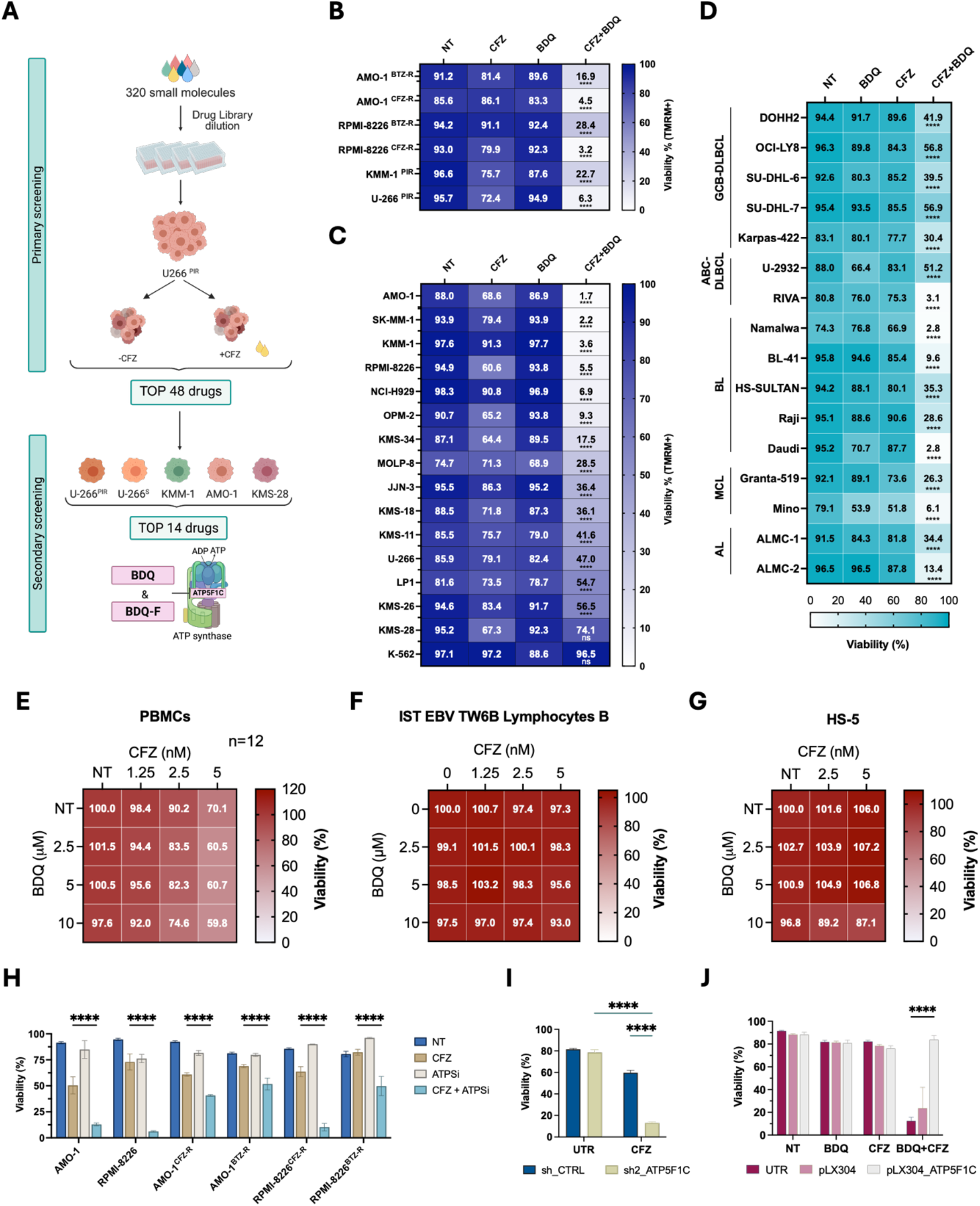
Bedaquiline Synergistically Enhances Proteasome Inhibitor Efficacy in Multiple Myeloma and Other Hematological Malignancies Through ATP5F1C Modulation. **(A)** Schematic overview of the functional drug screening strategy used to identify compounds that potentiate carfilzomib (CFZ) activity. A library of 320 small-molecule inhibitors was screened in U-266 PI-resistant (PIR) MM cells, with or without sublethal doses of CFZ. Top hits were validated in a secondary screen across five MM cell lines, identifying bedaquiline (BDQ) and its fumarate salt (BDQ-F) as strong synergistic partners of CFZ. **(B–C)** Cell viability assays showing the synergistic effect of BDQ and CFZ in PI-resistant (B) and PI-sensitive (C) MM cell lines. **(D)** Heatmap summarizing cell viability in systemic light chain amyloidosis (AL) and B-cell non-Hodgkin lymphoma (NHL) cell lines, including germinal center B-cell–like (GCB) and activated B-cell–like (ABC) diffuse large B-cell lymphoma (DLBCL), Burkitt’s lymphoma (BL), and mantle cell lymphoma (MCL), following treatment with BDQ, CFZ, or their combination. Viability was measured by PI staining at 96 hours in AL cell lines, and by TMRM staining at 72 hours in B-cell NHL lines. Compound concentrations are provided in Suppl. Tables S3–S4. **(E–G)** Dose–response matrices showing cell viability after BDQ-CFZ co-treatment in normal human cell models: **(E)** peripheral blood mononuclear cells (PBMCs) from 12 healthy donors; **(F)** EBV-transformed B lymphocytes (IST EBV TW6B; n = 3); **(G)** HS-5 bone marrow stromal cells (n = 3). Viability was measured by PI staining (PBMCs) or TMRM staining (IST EBV TW6B, HS-5) 96 hours post-treatment. **(H)** Viability of PI-sensitive (AMO-1, RPMI-8226) and PI-resistant (AMO-1^CFZ-R^, AMO-1^BTZ-R^, RPMI-8226^CFZ-R^, and RPMI-8226^BTZ-R^) MM cells treated with ATPSi, CFZ, or their combination. Viability was assessed 72 hours post-treatment using TMRM staining and flow cytometry. **(I)** Viability of AMO-1 cells expressing shCTRL or sh2_ATP5F1C treated with 3 nM CFZ or vehicle (DMSO), measured by TMRM staining 72 hours post-treatment. **(J)** Viability of AMO-1 cells transduced with pLX304 or pLX304_ATP5F1C and treated with 2.5 µM BDQ, 2.5 nM CFZ, the combination, or vehicle (DMSO). Viability was assessed 6 days post-treatment by TMRM staining and flow cytometry. Data represent the mean ± SD from three independent experiments unless otherwise indicated. Statistical significance was assessed using one-way ANOVA (****p < 0.0001).

To confirm that BDQ-CFZ synergy is mediated via ATP synthase inhibition, we tested the alternative oligomycin-derived ATP synthase inhibitor 1 (ATPSi). ATPSi-CFZ co-treatment significantly reduced viability of AMO-1 and RPMI-8226 cells, including their PI-resistant derivatives (p < 0.0001; Fig. 1H). Accordingly, ATP5F1C knockdown sensitized AMO-1 cells to CFZ (p < 0.0001; Fig. 1I; Suppl. Fig. 2A–B), Conversely, ATP5F1C overexpression mitigated BDQ-CFZ-induced cytotoxicity (p < 0.0001; Fig. 1J; Suppl. Fig. 2C-D). Collectively, these results demonstrate that BDQ and BDQ-F synergize with CFZ across diverse hematological malignancies, selectively inducing cytotoxicity via modulation of ATP5F1C, without harming healthy cells.

**Figure 2.**
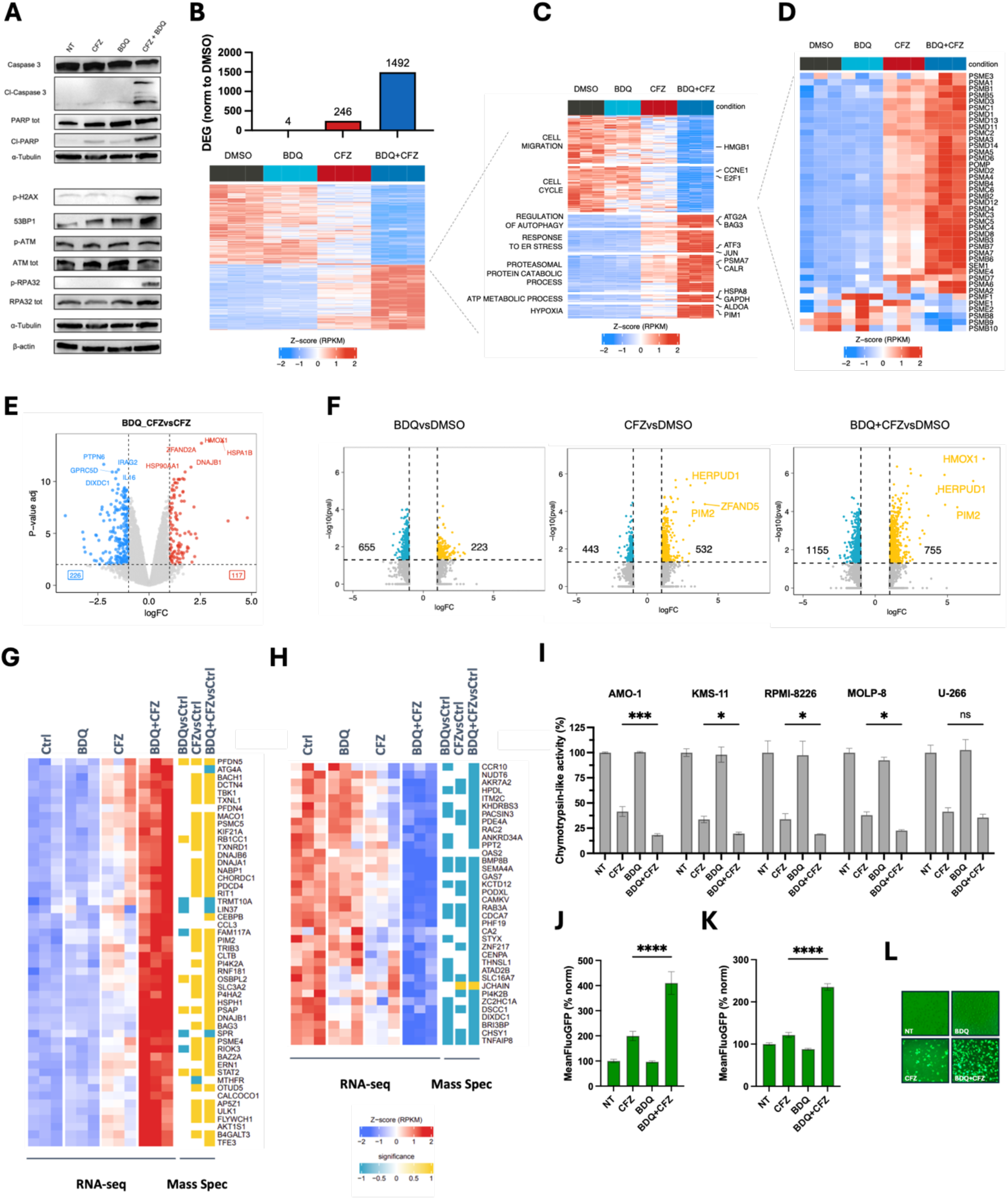
BDQ Potentiates Proteasome Inhibition and Specifically Synergizes with Proteasome Inhibitors in Multiple Myeloma. **(A)** Immunoblot analysis of apoptotic and DNA damage markers in AMO-1 cells treated for 16 hours with BDQ (5 µM), CFZ (2.5 nM), their combination (BDQ-CFZ), or left untreated (NT). α-Tubulin and β-actin were used as loading controls. **(B)** RNA sequencing analysis of AMO-1 cells reveals extensive transcriptional remodeling following BDQ-CFZ treatment. Bar plot: number of differentially expressed genes (DEGs) relative to untreated controls. Heatmap: hierarchical clustering of DEGs across treatment groups. **(C)** Gene ontology (GO) enrichment analysis of DEGs in BDQ-CFZ-treated cells, identifying upregulation of pathways involved in autophagy, endoplasmic reticulum (ER) stress, hypoxia, proteasomal degradation, ATP metabolism, and oxidative phosphorylation. **(D)** Heatmap of KEGG-defined proteasome gene expression showing transcriptional upregulation of proteasome components in BDQ-CFZ-treated cells. **(E)** Volcano plot of transcriptional changes comparing BDQ-CFZ vs. CFZ treatment. Red: significantly upregulated genes; green: significantly downregulated genes; gray: non-significant. **(F)** Volcano plots of label-free quantitative proteomics comparing protein expression changes in AMO-1 cells treated with BDQ, CFZ, or the combination (BDQ-CFZ). Yellow: upregulated; light blue: downregulated. **(G-H)** Integrated heatmaps of RNA-seq and proteomics data showing **(G)** upregulated and **(H)** downregulated transcripts and proteins, highlighting convergence on cellular stress response pathways. **(I)** Chymotrypsin-like proteasome activity was measured 5 hours after treatment with BDQ (5 µM or 10 µM for U-266), CFZ (2.5 nM), or their combination in a panel of MM cell lines (AMO-1, KMS-11, RPMI-8226, MOLP-8, and U-266). BDQ significantly enhanced CFZ-mediated proteasome inhibition in all cell lines except U-266, which showed reduced sensitivity to the combination. **(J-K)** Quantification of proteasome inhibition using a Ub-G76V-GFP reporter assay in AMO-1 and AMO-1^BTZ-R^ cells. Flow cytometry analysis shows increased GFP accumulation upon BDQ-CFZ treatment, indicating enhanced proteasome inhibition. Fluorescence intensity is normalized to untreated controls and represents the average of three independent experiments (mean ± SD). Statistical analysis confirms significant synergy in both parental and resistant cells (****p < 0.0001, one-way ANOVA). **(L)** Representative fluorescence microscopy images of Ub-G76V-GFP–expressing AMO-1 cells treated with BDQ, CFZ, or the combination, showing marked GFP accumulation in BDQ-CFZ–treated cells.

To elucidate the basis of BDQ-CFZ synergy in MM, we first profiled early apoptotic and stress responses. Immunoblotting of AMO-1 cells 16 hours post-treatment revealed robust activation of caspase-3, PARP cleavage, and increased phosphorylation of H2A.X and RPA32, indicating enhanced apoptosis and DNA damage (Fig. 2A). BDQ-CFZ combination further reduced oxygen consumption rate and elevated mitochondrial ROS levels as compared to single treatments (Suppl. Fig. 2E-F). Transcriptomic profiling at 12 hours post-treatment, prior to detectable viability loss (Suppl. Fig. 3A–C), revealed 246 differentially expressed genes in response to CFZ, whereas BDQ alone had minimal transcriptional effects. Strikingly, BDQ-CFZ co-treatment led to 1,492 differentially expressed genes (Fig. 2B). These included upregulation of autophagy, proteasome catabolism, ER stress, and oxidative phosphorylation pathways, alongside downregulation of cell cycle and migration-related genes (Fig. 2C). Notably, BDQ enhanced CFZ-induced expression of proteasome subunits (Fig. 2D), consistent with previously described compensatory feedback mechanisms.^21^ Independent RT-qPCR analysis confirmed several of the most prominently upregulated and downregulated genes (Fig. 2E; Suppl. Fig. 3D–F). Complementary label-free quantitative proteomics further substantiated the synergistic impact of BDQ-CFZ treatment, revealing a marked increase in upregulated proteins compared to single-agent exposure (Fig. 2F). Integrated analysis of transcriptomic and proteomic data highlighted a convergent activation of cellular stress response pathways (Fig. 2G-H).

**Figure 3.**
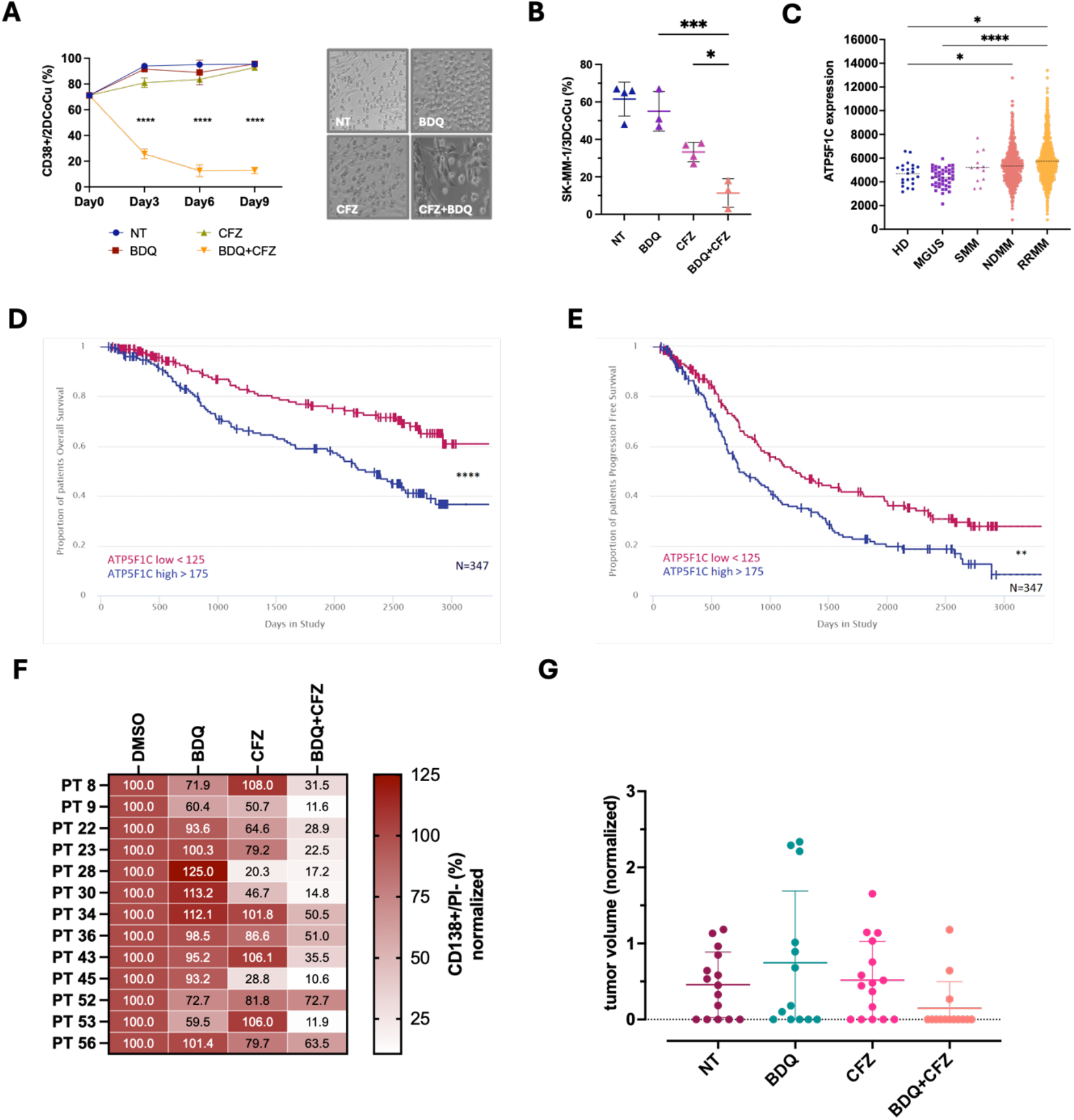
BDQ-CFZ Potentiates Proteasome Inhibition in Patient-Derived and In Vivo Models. **(A–B)** BDQ-CFZ co-treatment potentiates MM cell death in bone marrow–mimicking microenvironments. **(A)** AMO-1 cells were cultured in 2D with GFP+ HS-5 stromal cells and treated with 5 µM BDQ and 2.5 nM CFZ at day 0 and day 3. MM cell viability was assessed by flow cytometry using CD138-APC staining at the indicated time points and normalized to GFP+ stromal cells (left). Representative bright-field image (right) shows AMO-1 cells on HS-5 stromal monolayer 3 days post-treatment. Statistical analysis: two-way ANOVA (n = 2–3). **(B)** In a 3D co-culture system, GFP+ HS-5 stromal cells and tRFP+ SK-MM1 MM cells were seeded into scaffolds and treated with CFZ (6 nM), BDQ (10 µM), or their combination. MM cell viability was quantified by flow cytometry 6 days post-treatment. BDQ-CFZ co-treatment significantly reduced the viability of MM cells compared to single agents (Tukey’s multiple comparisons test, p = 0.0242; n = 3–4). *P < 0.05; **P < 0.01; ***P < 0.001. **(C)** ATP5F1C expression increases with disease progression across healthy donors (HD), MGUS, SMM, NDMM, and RRMM groups (data from GSE2658, GSE5900, GSE31161). **(D-E)** High ATP5F1C expression is associated with poor prognosis in MM patients, based on progression-free and overall survival analyses from the CoMMpass study (IA22, n = 347; ****P< 0.0001). Statistical significance was determined by two-way ANOVA (****p < 0.0001; *p < 0.05). **(F)** BDQ enhances CFZ-induced cytotoxicity in primary MM cells. BMMCs from 13 MM patients were treated with CFZ, BDQ, or the combination for 96 hours. Viability was assessed by flow cytometry using PI and CD138 stainings and normalized to DMSO controls. Drug concentrations for each patient are listed in Suppl. Table S5. **(G)** In vivo validation using a zebrafish xenograft model. RPMI-8226 cells (tRFP+) were injected into zebrafish embryos and treated with BDQ, CFZ, or their combination. Tumor burden at 72 hours post injection (hpi) was normalized to tumor area at 24 hpi (DMSO: n = 15; CFZ: n = 16; BDQ: n = 13; BDQ-CFZ: n = 14). Abbreviations: PBMCs, peripheral blood mononuclear cells; BMMCs, bone marrow mononuclear cells; s.e.m., standard error of the mean; hpi, hours post-injection; HD, healthy donors; MGUS, monoclonal gammopathy of undetermined significance; SMM, smoldering MM; NDMM, newly diagnosed MM; RRMM, relapsed/refractory MM.

Importantly, we observed that BDQ potentiated CFZ-mediated chymotrypsin-like proteasome inhibition across MM cell lines, but not in low-responder U-266 cells (Fig. 2I). This enhancement was selective to chymotrypsin-like activity, as caspase-like and trypsin-like activities remained largely unaffected (Suppl. Fig. 4A-B). Proteasome blockade was confirmed in AMO-1 and AMO-1^BTZR^ cells by expressing a short-lived GFP fusion protein (Ub-G76V-GFP), showing increased GFP signal following BDQ-CFZ treatment, before cell death commitment (Fig. 2J-L; Suppl. Fig. 4C-D). BDQ synergized with other PIs, including BTZ, IXA, and MRZ, as well as the ubiquitin-activating enzyme inhibitor TAK-243 across a broad range of PI-sensitive and PI-resistant MM cell lines (Suppl. Fig. 5). However, no synergy was observed with standard agents (e.g., dexamethasone, venetoclax, lenalidomide, melphalan, or doxorubicin), underscoring the specificity of BDQ’s interaction with ubiquitin-proteasome system (UPS) targeting drugs (Suppl. Fig. 4E). Together, these findings reveal that BDQ-CFZ co-treatment induces broad transcriptional and proteomic stress responses, selectively enhancing proteasome inhibition and cytotoxicity in MM cells.

To assess the translational potential of BDQ-CFZ, we tested its efficacy in 2D and 3D co-culture systems with HS-5 stromal cells, mimicking the bone marrow niche. The combination significantly increased MM cell death over time in both settings, while sparing stromal cells (Fig. 3A–B), demonstrating selective activity even in protective microenvironments. Public gene expression data highlighted that ATP5F1C is upregulated in NDMM and RRMM vs. healthy donors, and more highly expressed in RRMM than in MGUS (Fig. 3C). Analysis of patient datasets (CoMMpass IA22, n=347) revealed that high ATP5F1C expression, BDQ’s mitochondrial target, correlates with poor progression-free and overall survival (p < 0.01; Fig. 3D-E). Ex vivo treatment of CD138^+^ cells from 13 MM patient bone marrow samples showed that BDQ significantly enhanced CFZ-induced cytotoxicity, regardless of cytogenetic background or treatment history (p < 0.01; Fig. 3F; Suppl. Table S2). Finally, in a zebrafish xenograft model, BDQ-CFZ reduced DsRed^+^ tumor burden (Fig. 3G), confirming in vivo efficacy. These findings support the safety, specificity, and clinical relevance of BDQ-CFZ therapy for MM, and highlight ATP5F1C as a potential biomarker for therapeutic response.

The present study demonstrates that the ATP synthase inhibitor BDQ, along with its fumarate salt form (BDQ-F), exhibits consistent and robust synergy with agents targeting the UPS across various MM models, as well as in amyloidosis and diverse B-cell malignancies. BDQ is an orally bioavailable antimicrobial agent originally developed to inhibit ATP synthase in *Mycobacterium tuberculosis*, and has revolutionized the therapeutic landscape for multidrug-resistant tuberculosis.^22^ In addition, BDQ can inhibit the γ-subunit of human mitochondrial ATP synthase, ATP5F1C.^23^ Importantly, ATP5F1C plays an essential role in mitochondrial ATP production and has been linked to increased tumor aggressiveness and adverse clinical outcomes.^24,25^ Mechanistic validation using both genetic knockdown and pharmacologic inhibition of ATP5F1C phenocopied BDQ-CFZ cytotoxicity, while overexpression conferred resistance, thereby confirming ATP5F1C as the key functional target. Transcriptomic and proteomic analyses revealed that BDQ co-treatment with CFZ amplified stress-response pathways, inducing DNA damage, apoptosis, and proteotoxic stress. Importantly, BDQ potentiated CFZ-mediated inhibition of chymotrypsin-like proteasome activity. This suggests that BDQ, by impairing ATP production and disrupting oxidative phosphorylation, exacerbates proteotoxic stress induced by proteasome inhibition. The observed synergy highlights a therapeutic opportunity to exploit metabolic vulnerabilities as a means to overcome PIs resistance. By acting through energy stress pathways, BDQ circumvents canonical resistance mechanisms and specifically sensitizes cells to UPS-targeted therapy.

Although BDQ’s clinical approval facilitates translation potential, its preferential inhibition of mycobacterial ATP synthase at submicromolar concentrations^23^ raises critical considerations. The effective in vivo dosing required to inhibit ATP5F1C in human cancer cells must be rigorously defined, and more selective inhibitors targeting the γ subunit of mitochondrial ATP synthase should be developed. It also remains to be determined whether inhibition of other subunits (e.g., α or β) would confer similar antitumor synergy. However, unlike ATP5F1C, expression of ATP5F1A and ATP5F1B did not correlate with inferior prognosis in MM patients (Suppl. Fig. 6), further supporting the specificity to target ATP synthase γ. Besides, ATP depletion may disrupt other mitochondrial functions or intersect with the tricarboxylic acid cycle, contributing to cytotoxicity. Dissecting these downstream consequences will be essential to fully elucidate the mechanism of action and to rationally design future combination regimens.

In summary, this study supports a therapeutic strategy that combines proteasomal and metabolic stress to restore drug sensitivity in MM. The clinical availability of BDQ offers an immediate translational path, while future work should focus on refining target specificity, optimizing dosing, and extending this approach to other malignancies marked by proteotoxic stress.

## ETHICS APPROVAL AND CONSENT TO PARTICIPATE

PBMCs from healthy donors and bone marrow aspirates from MM patients were obtained from the local Blood Bank and the Hematology Unit at Città della Salute e della Scienza Hospital (Turin, Italy). Informed consent was obtained from all participants following the Declaration of Helsinki, and the study was approved by the local ethics committee (protocol no. 00143/2022). All animal experiments were conducted in compliance with institutional guidelines and approved by the Ethical Committee of the University of Torino.

## AUTHORS’ CONTRIBUTIONS

M.C., carried out most of the experiments and contributed to the interpretation of biological data with E.M., M.L., M.P., and M.M.; E.M. performed drug screening experiments and analysis; B.L. studied BDQ combinations with proteasome inhibitors; A.T. and F.A., performed RNA-Seq experiments and bioinformatics analyses; S.O. supervised bioinformatics analyses; P.E.P. supervised biochemical assays; A.L., M.D.A., F.G. and B.B. provided clinically annotated MM samples; M.C., M.M. and C.B. performed zebrafish xenograft experiments; Eu.M. and N.M. participated in the experimental design and the interpretation of biological data; R.P. designed the studies and supervised the project; M.C. and R.P. wrote the manuscript; all authors were involved in the final version of the manuscript.

## ACKNOWLEDGEMENTS

The authors gratefully acknowledge the Advanced Microscopy Open Lab (OLMA@MBC) and Dr. Marta Gai for assistance with image acquisition of zebrafish xenotransplanted embryos; Dr. Lorenzo Prever, and Profs. Giorgio R. Merlo and Emilio Hirsch for their guidance with zebrafish xenograft experiments; Drs. Karel Harant and Pavel Talacko at the Laboratory of Mass Spectrometry (BIOCEV, Vestec, Czech Republic) for performing and analyzing the label-free quantitative proteomic experiments; and Drs. Ivan Zaggia, Marcello Turi, and Anna Maria Gullà for their technical and scientific support with metabolic and immunological assays.

