## Supplemental Materials for "Bedaquiline Amplifies Proteasome Inhibitor Efficacy and Overcomes Resistance in Multiple Myeloma"

[1] Department of Molecular Biotechnology and Health Sciences, University of Turin, Turin, Italy; [2] Molecular Biotechnology Center (MBC) "Guido Tarone", Turin, Italy; [3] Dana Farber Cancer Institute, Boston, MA, United States; [4] Città Della Salute e della Scienza Hospital, Turin, Italy; [5] Department of Life Sciences and Systems Biology, University of Turin, Turin, Italy; [6] Candiolo Cancer Institute, FPO-IRCCS, Turin, Italy.

### SUPPLEMENTAL MATERIALS AND METHODS

#### Cell Culture Conditions and Reagents

Human multiple myeloma (MM) cell lines—KMM-1, U-266, KMM-1$^{PIR}$, U-266$^{PIR}$, KMS-11, NCI-H929, LP-1, AMO-1, OPM-2, KMS-18, KMS-26, KMS-28, KMS-34, RPMI-8226, SK-MM-1, MOLP-8, and JJN-3—along with the murine MM cell line 5TGM1, the chronic myelogenous leukemia cell line K-562, the human stromal cell line HS-5, the Burkitt lymphoma (BL) cell lines (RAJI, HS-Sultan, Daudi, BL-41, and Namalwa), the mantle cell lymphoma (MCL) cell lines (Granta-519 and Mino), and the diffuse large B-cell lymphoma (DLBCL) cell lines including RIVA and U-2932 (ABC-DLBCL), and OCI-LY8, SU-DHL-6, SU-DHL-7, DoHH2, and KARPAS-422 (GCB-DLBCL) were obtained from the DSMZ (Braunschweig, Germany), the ATCC (Manassas, VA, USA), or generated in-house. All cell lines were authenticated by DNA fingerprinting using the GenePrint system (Promega, Madison, WI, USA). Drug-resistant MM lines AMO-1$^{CFZ\text{-}R}$, AMO-1$^{BTZ\text{-}R}$, RPMI-8226$^{CFZ\text{-}R}$, and RPMI-8226$^{BTZ\text{-}R}$ were kindly provided by Dr. Lenka Besse (St. Gallen, Switzerland).[1,2] EBV-immortalized B-lymphocytes (IST-EBV-TW6B) were generously provided by Prof. Valentina Cauda (Politecnico di Torino, Italy). HS-5 (GFP+) cells and RPMI-8226 (tRFP+)-positive cells were derived by transduction with pLKO-GFP and pLemir control vectors, respectively. All cell lines, unless otherwise specified, were cultured in RPMI 1640 (EuroClone, Pero, Italy) supplemented with 10% fetal bovine serum (FBS; Sigma-Aldrich, St. Louis, MO, USA), 2 mM L-glutamine, 100 U/mL penicillin, and 100 μg/mL streptomycin (Gibco). IST-EBV-TW6B cells were maintained in RPMI 1640 with 20% FBS and 1% non-essential amino acids. HEK 293T cells (DSMZ) were cultured in DMEM with 10% FBS and the same supplementation. The AL amyloidosis cell lines ALMC-1 and ALMC-2 were maintained in Iscove’s Modified Dulbecco’s Medium (Gibco, Thermo Fisher Scientific) supplemented with 10% FBS (Biological Industries), 2 mM L-glutamine, 1% penicillin-streptomycin, 1 ng/mL human interleukin-6 (PeproTech), and 10 ng/ml IGF1. All cultures were maintained at 37°C in a humidified incubator with 5% $CO_2$. Carfilzomib (CFZ), bortezomib (BTZ), ixazomib (IXA), melphalan (MEL), doxorubicin (DOX), dexamethasone (DEXA), lenalidomide (LENA), venetoclax (VEN), bedaquiline (BDQ), BDQ-fumarate (BDQ-F), and ATP synthase inhibitor 1 (ATPSi) were purchased from Selleckchem (Munich, Germany) or MedChemExpress (Stockholm, Sweden). TAK-243 (TAK) was obtained from TargetMol (Boston, MA, USA).

**Drug screening**

Primary compound screening was performed using a selective small-molecule inhibitor library (L3500, Selleck Chemicals) consisting of 320 inhibitors targeting 123 key proteins involved in diverse signaling pathways. U-266$^{PIR}$ multiple myeloma (MM) cells were treated with each compound at four concentrations (10 μM, 1 μM, 100 nM, and 10 nM). After 2 hours, cells were exposed to either a sublethal dose of carfilzomib (CFZ; 20 nM) or dimethyl sulfoxide (DMSO) as a vehicle control. Cell viability was measured at baseline (0 h) and 72 hours post-treatment using the CellTiter-Glo Luminescent Cell Viability Assay (Promega), and results were collected in duplicate. Growth rate (GR) was calculated by normalizing the luminescent signal at 72 hours to the baseline signal and further normalized to the DMSO-treated controls. Drug combination effects were quantified using the Excess Over Bliss (EOB) model, calculated from GR values using the following formula:

1 − GR(combination)] – [1 − GR(CFZ)] – [1−GR(drug)] + [1−GR(CFZ)][1 − GR(drug)][3]

Compounds yielding EOB ≥ 0.4 at any concentration were considered potential hits and shortlisted for secondary validation in a panel of five MM cell lines (U-266$^{PIR}$, U-266, KMM-1, AMO-1, and KMS-28). Compounds that achieved EOB ≥ 0.4 in at least three cell lines were designated top synergistic candidates (TOP14) and listed in Suppl. Table S1.

**Virus Production and In Vitro Transduction**

High-titer lentiviral stocks were generated in 293T cells by co-transfecting the expression vector (pLKO, pLX304, or pLX301) with packaging vectors (pCMVdR8.74 and VSV-G/pMD2.G), using the Effectene Transfection Reagent (Qiagen, Milan, Italy) according to the manufacturer's protocol. Viral supernatants were collected over 36–60 hours, filtered through a 0.22 μm filter, and concentrated using the Lenti-X Concentrator (ClonTech). Concentrated viruses were resuspended in cold phosphate-buffered saline (PBS) and stored at –80°C. Cells (1 × $10^5$/mL) were transduced in the presence of 8 μg/mL polybrene. Fresh medium was added 2 hours post-infection. Stable cell lines were selected using 1 μg/mL puromycin or blasticidin (Sigma-Aldrich). Cell viability was assessed at 48 or 72 hours post-transduction (hpt) using tetramethylrhodamine methyl ester (TMRM; Molecular Probes, Eugene, OR, USA) and flow cytometry.

**Gene Silencing and Overexpression of ATP5F1C**

Lentiviral vectors encoding short hairpin RNAs (shRNAs) targeting ATP5F1C (shATP5F1C) or a non-targeting control (shCTRL) were generated by subcloning the respective sequences into the pLKO.1-puro vector (Addgene #8453) using AgeI–EcoRI restriction sites. shRNA sequences are listed in Suppl. Table S6. AMO-1 cells were transduced with shATP5F1C or shCTRL lentiviruses, selected with puromycin, and silencing efficiency was confirmed by RT-qPCR and immunoblotting.

For overexpression, the ATP5F1C open reading frame was obtained from the DNASU plasmid repository (pLX304-ATP5F1C, HsCD00441174) and used to produce lentiviruses. AMO-1 cells were transduced with pLX304-ATP5F1C or empty vector (pLX304), followed by blasticidin selection. Overexpression was validated by RT-qPCR and immunoblotting.

**Ub-G76V-GFP Proteasome Inhibition Reporter**

The pLX301_Ub-G76V-GFP vector was derived from pEGFP-N1-Ub-G76V-GFP (Addgene #11941, gift from Nico Dantuma), as described.[4] For proteasome inhibition assays, MM cell lines AMO-1_UbGFP and AMO-1_UbGFP$^{BTZ-R}$ were seeded at $1x10^5$ cells/mL. AMO-1_UbGFP and AMO-1_UbGFP$^{BTZ-R}$ cells were treated with 2.5 nM and 12.5 nM CFZ, respectively, 10 µM BDQ, or their combination. Cell viability was assessed by TMRM staining, and GFP-positive cells along with their fluorescence intensity were analyzed by flow cytometry and fluorescence microscopy using a ZEISS Axio Observer inverted microscope (10× objective) at 10, and 24 hpt.

**ATP-Lite Assay**

Cell viability/proliferation was measured using the CellTiter-Glo® Luminescent Cell Viability Assay (Promega). Cells were seeded at $1 \times 10^5$/mL and analyzed at day 0 and at the indicated timepoints. Ten microliters of cells were mixed with an equal volume of CellTiter-Glo® reagent in a white 384-well plate, in technical duplicate. Plates were shaken in the dark for 2 minutes and incubated at room temperature (RT) for 10 minutes. Luminescence was measured using the Synergy™ 2 Multi-Mode Microplate Reader (BioTek Instruments, Winooski, VT, USA).

**CCK-8 Assay**

Cell viability/proliferation was also assessed using the Cell Counting Kit-8 (CCK-8; Dojindo). Cells were plated at $1 \times 10^5$/mL and evaluated at day 3 or day 4. 90 µL of cell suspension were mixed with 10 µL of CCK-8 reagent in a 96-well plate, in technical quadruplicate. Plates were incubated at 37°C for 1–4 hours (without $CO_2$). Absorbance at 450 nm was measured using the Synergy™ 2 reader.

**Cell Cycle and Cell Viability Assays by Flow Cytometry**

Apoptosis was measured by flow cytometry after staining with tetrametylrodamine methyl ester (TMRM; Molecular Probes, Eugene, Oregon, USA) or Annexin V-FITC Kit (Miltenyi Biotec, Bergisch Gladbach, Germany), according to the manufacturer's instructions. $CD138^+$ cells were identified using anti-human CD138-APC (clone 44F9; Miltenyi). Cell cycle analysis was performed by PI staining. Cells were washed in PBS, incubated with RNase (0.14 mg/mL) and stained with PI (28.57 μg/mL). Data were acquired using BD FACSCelesta™ or BD Accuri™ instruments and analyzed with FACSDiva v8.0 (BD Biosciences).

**Proteasome Activity Assay**

Chymotrypsin-like, trypsin-like, and caspase-like proteasome activities were measured using the Proteasome-Glo™ Cell-Based Assay (Promega, G1180) according to the manufacturer's protocol. MM cell lines AMO-1, KMS-11, RPMI-8226, MOLP-8, U-266 cell lines were plated at $1\times10^5$ cells/mL and treated with 2.5 nM CFZ, 5 μM BDQ, or their combination. Proteasome activity was assessed 5 hours post-treatment in technical quadruplicates. For each condition, 50 μL of cell suspension was combined with 50 μL of the appropriate Proteasome-Glo™ reagent (Chymotrypsin-like, Trypsin-like, or Caspase-like) in white 96-well plates. Plates were shaken at 700 rpm for 2 minutes and incubated for 10 minutes at room temperature in the dark. Luminescence was recorded using a Synergy™ 2 Multi-Mode Microplate Reader (BioTek Instruments, Winooski, VT).

**Mitochondrial Superoxide Detection**

Mitochondrial superoxide was measured using the MitoSOX™ Red Indicator (Thermo Fisher Scientific), according to the manufacturer's protocol, and analyzed by flow cytometry.

**Measurement of Oxygen Consumption Rate**

Oxygen consumption rate (OCR) was monitored over time using the Resipher™ system (Lucid Scientific, Atlanta, GA, USA). AMO-1 cells were seeded at 20,000 cells/well in 96-well plates and incubated at 37 °C, 5% $CO_2$. OCR was recorded via the Lucid Lab web app, with data captured every 60 minutes, and monitored for 36 h after treatment with CFZ and BDQ at the indicated concentrations. Measurements are the average of five independent replicates.

**Western Blotting**

Cells were lysed in buffer (20 mM Tris-HCl pH 7.4, 150 mM NaCl, 5 mM EDTA, 1% Triton X-100, 1 mM PMSF, 10 mM NaF, 1 mM $Na_3VO_4$, and protease inhibitors; Roche). Lysates were incubated at 4°C for 30 min, centrifuged at 13,000 × g for 15 min, and protein quantified by Bio-Rad DC assay. Equal amounts of protein were resolved by SDS-PAGE, transferred to nitrocellulose membranes, blocked in 5% milk in PBS-T for 1 h, then incubated overnight with primary antibodies in 5% BSA + $NaN_3$. Membranes were washed and incubated with secondary antibodies for 1 h, then visualized with Immobilon Western HRP substrate (Merck). Antibodies used are listed in Supplementary Table S6.

**RNA Extraction and Reverse Transcription-quantitative PCR (RT-qPCR)**

Total RNA was isolated using the RNeasy Mini Kit (Qiagen). RNA was treated with RQ1 RNase-free DNase (Promega) and reverse-transcribed using the OneScript Plus cDNA Synthesis Kit (ABM, Canada) or SuperScript III (Invitrogen, Carlsbad, CA, USA). RT-qPCR was performed in 384-well plates on a Bio-Rad iCycler using iQ SYBR Green or BlasTaq™ 2X qPCR MasterMix (ABM), following manufacturer instructions. Cycling: 95°C for 10 min, then 40 cycles of 95°C for 15 s and 60°C for 1 min. Primer sequences (designed via PrimerBLAST) are available upon request and listed in Supplementary Table S2. Specificity was validated by melting curve, standard curve slope, and linearity. GAPDH served as the endogenous control. Expression levels were normalized to control infections (scrambled shRNA, empty vector, or uninfected).

**RNA-sequencing**

AMO-1 cells were treated with DMSO, CFZ (2.5 nM), BDQ (5 μM), or their combination, and harvested at 6, 12, and 18 hpt. RNA was extracted using the RNeasy Mini Kit and quality assessed via Agilent TapeStation. Libraries were prepared with the TruSeq Stranded mRNA Kit (Illumina) and sequenced on the Illumina NextSeq 1000. Reads were aligned to hg38 using STAR v2.7.1a. Gene counts were obtained with featureCounts v1.6.3 using GENCODE v42.[5] Differential expression was determined by GLM and likelihood ratio tests; genes with |logFC| >1 and FDR <0.01 were considered significant. Gene set enrichment was performed using the SnakeGSEA pipeline with multiGSEA (https://github.com/molinerisLab/SnakeGSEA).[6] Heatmaps were generated using the ComplexHeatmap R package.[7]

**Mass Spectrometry (MS) and Proteomic Data Analysis**

Cells were lysed by boiling at 95°C for 10 minutes in 100 mM triethylammonium bicarbonate (TEAB) containing 2% sodium deoxycholate (SDC), 40 mM chloroacetamide, and 10 mM tris(2-carboxyethyl)phosphine (TCEP), followed by sonication (Bandelin Sonoplus Mini 20, MS 1.5). Protein concentration was determined using the BCA Protein Assay Kit (Thermo Scientific), and 30 μg of protein per sample was used for MS preparation. The volume was adjusted to 50 μL with 100 mM TEAB containing 2% SDC. Samples were processed using the SP3 protocol, as described by Hughes et al. Briefly, 5 μL of SP3 beads were added to 30 μg of protein, and the final volume was brought to 50 μL with 100 mM TEAB. Ethanol was added to a final concentration of 60% (v/v) to induce protein binding. Samples were mixed and incubated for 5 minutes at room temperature (RT). After magnetic separation, the supernatant was discarded, and the beads were washed twice with 180 μL of 80%

ethanol. Proteins were digested on-bead overnight at 37°C using trypsin at a 1:30 enzyme-to-protein ratio, in 100 mM TEAB. Following digestion, samples were acidified to 1% trifluoroacetic acid (TFA) and desalted using in-house packed C18 stage tips (Empore disks).[8] Peptides were analyzed by nanoLC-MS/MS using an Aurora Ultimate TS column (25 cm × 75 µm ID, 1.7 µm particles; Ion Opticks). Mobile phase A consisted of water with 0.1% formic acid, and mobile phase B consisted of acetonitrile with 0.1% formic acid. Samples were first loaded onto a trap column (C18 PepMap100, 5 µm, 300 µm × 5 mm; Thermo Scientific) for 4 minutes at 18 µL/min using a loading buffer of water, 2% acetonitrile, and 0.1% TFA. Peptides were then eluted with a linear gradient from 4% to 35% B over 60 minutes. Eluted peptides were ionized by electrospray and analyzed on a Thermo Scientific Orbitrap Ascend in data-independent acquisition (DIA) mode. Survey scans were acquired from 350–1400 m/z at 60,000 resolution (at 200 m/z) with an ion target of $4 \times 10^5$. DIA scans were collected at 30,000 resolution with an AGC target of 1000% and maximum injection time set to "Auto." The 400–1000 m/z range was segmented into 30 windows of 20 Da each. HCD fragmentation was performed at a normalized collision energy of 28%. Data were processed and quantified using Spectronaut 18 in directDIA mode.[9] Searches were conducted against the human UniProt database (June 2023 release; 20,605 entries). Enzyme specificity was set to C-terminal cleavage after Arg and Lys, allowing cleavage at proline residues and up to two missed cleavages. Carbamidomethylation of cysteines was defined as a fixed modification; N-terminal acetylation and methionine oxidation were set as variable modifications. False discovery rate (FDR) was controlled at 1% for peptide-spectrum matches (PSMs), peptides, and proteins. Quantification was based on MS2-level data. Precursor and protein posterior error probability (PEP) thresholds were set to 0.01 and 0.05, respectively. Processed data were exported and further analyzed using Perseus v1.6.15.0.[10]

**2D Co-culture**

HS-5 ($GFP^+$) and AMO-1 cells were seeded in 24-well plates at $5 \times 10^4$/mL and $1 \times 10^5$/mL, respectively. Cells were treated with CFZ (2.5 nM), BDQ (5 µM), or the combination at day 0 and 3. Every 3 days, cells were collected and replated. Cell viability was measured every 3 days over 9 days using TMRM staining and flow cytometry.

**3D Co-culture**

3D scaffolds were prepared following the protocol by Belloni et al.,[11] with adaptations for multiple myeloma (MM) cell culture under static conditions. Briefly, scaffold discs were created from Spongostan™ sheets (Ethicon, Inc., Somerville, NJ, USA) using a sterile 4 mm biopsy punch. Each disc was pre-seeded with $2x10^5$ GFP-positive HS-5 stromal cells. After 24 hours, $5x10^5$ tRFP-positive SK-MM-1 cells were added to each scaffold. The scaffolds were then cultured in 1 mL of RPMI-1640 medium in 24-well plates. To minimize the presence of non-adherent cells, the scaffolds were transferred to new wells every 24 hours for three days. Drug treatments were administered 72 hours after seeding the SK-MM-1 cells, using sublethal doses of CFZ, BDQ, or a combination of the two drugs. Six days post-treatment, cells were released from the scaffolds using Liberase™ (25 µg/mL, Merck Millipore, Burlington, MA, USA) and analyzed by flow cytometry to assess the ratio of stromal and tumor cells.

**Gene Expression Data**

The expression and prognostic value of ATP5F1C messenger RNA (mRNA) in terms of Overall Survival (OS) and Progression Free Survival (PFS) were determined using publicly available RNA-sequencing (RNAseq) data from newly diagnosed MM patients from the Multiple Myeloma Research Foundation's (MMRF) CoMMpass study (https://research.themmrf. org/; release IA22). Data were analyzed with GraphPad prism 9.1 to retrieve mean ATP5F1C expression (Transcript per million, TPM) and standard deviation.

**MM Patient and Healthy Donor Samples**

Peripheral blood mononuclear cells (PBMCs) from healthy donors and bone marrow (BM) aspirates from MM patients were obtained from the local Blood Bank (Città della Salute e della Scienza Hospital, Turin). PBMCs were isolated from buffy coats via Ficoll-Hypaque density gradient, resuspended in RPMI + 10% FBS, and seeded at $5\times10^5$ cells/mL. BM mononuclear cells were isolated by Ficoll gradient centrifugation following Miltenyi's protocol. $CD138^+$ plasma cells were identified by flow cytometry using anti-CD138-APC (clone 44F9, Miltenyi), and samples with ≥10% $CD138^+$ cells were used for downstream assays. Cells were seeded at $2.5\times10^5$ cells/mL in RPMI + 10% FBS and treated as described in Suppl. Table S3.

**Zebrafish Xenograft Assay**

Approximately 250 RPMI-8226 cells stably expressing DsRed fluorescent protein were injected (2 nL) into the yolk sac of 48 hours post-fertilization (hpf) embryos with a manual microinjector (Eppendorf, Germany). Embryos were maintained at 28°C in embryo medium supplemented with 0.003% PTU, 1 g/L glucose, and 5 mM L-glutamine. At 24 hours post-injection (hpi), xenograft-positive embryos were identified by fluorescence microscopy (ZEISS Axio Observer, 10× objective), randomly allocated into four treatment groups (DMSO, 2.5 nM CFZ, 10 µM BDQ, or combination), and placed individually into 48-well plates. Tumor burden was assessed at 72 hpi by fluorescence microscopy and quantified using ImageJ as the fluorescence area normalized to the 24 hpi baseline. Adult zebrafish were euthanized with a tricaine overdose. All procedures were approved by the University of Torino Ethical Committee and the Italian Ministry of Health.

**Synergy Score**

Synergistic score of the combinatorial treatment with BDQ or BDQ-F and CFZ was assessed using SynergyFinder+. The score was calculated based on either the Bliss Score (BS) or the ZIP Score.

**Statistical analyses**

Statistical analyses were performed with GraphPad Prism 10 (GraphPad Software Inc.). Statistical significance of differences observed (in both *in vitro* and *in vivo* experiments) was determined by Student t test or one-way ANOVA; differences were considered significant when P value was <.05 (*), <.01 (**), <.001 (***), or <.0001 (****).

**SUPPLEMENTAL FIGURES AND LEGENDS**

**Figure S1. Synergistic Effects of Bedaquiline (BDQ) and Its Fumarate Salt (BDQ-F) with Carfilzomib (CFZ) in Multiple Myeloma Cell Lines. (A)** 3D dose-response matrices showing the combinatorial effects of BDQ and CFZ in AMO-1, RPMI-8226, KMS-11, SK-MM-1, KMM-1, and KMM-1$^{PIR}$ cell lines. Cell viability was assessed by TMRM staining and analyzed by flow cytometry 72 hours post-treatment. Bliss Synergy (BS) scores were calculated using the SynergyFinder+ tool (www.synergyfinder.org). **(B)** Dose-response matrices of BDQ-F and CFZ combinations across a panel of MM cell lines and the K-562 chronic myeloid leukemia cell line. Viability was measured using CellTiter-Glo® Luminescent assay 72 hours post-treatment and normalized to luminescence values at time 0. Synergy scores (SS) were calculated using the Bliss independence model.

**Figure S2. ATP5F1C Knockdown and Overexpression Efficiency; BDQ-CFZ Effects on Oxygen Consumption Rate and Mitochondrial ROS Levels (A)** ATP5F1C knockdown efficiency in AMO-1 cells transduced with shRNA targeting ATP5F1C (sh2_ATP5F1C) or scrambled control (shCTRL), quantified by RT-qPCR following puromycin selection. **(B)** Representative western blot showing ATP5F1C protein levels in AMO-1 cells expressing shCTRL or sh2_ATP5F1C. α-Tubulin in was used as a loading control. **(C)** Relative ATP5F1C expression in AMO-1 cells transduced with pLX304 vector or pLX304_ATP5F1C, assessed by RT-qPCR to confirm overexpression. **(D)** Representative western blot of ATP5F1C and V5-Tag in AMO-1 cells transduced with pLX304 vector or pLX304_ATP5F1C. Vinculin was used as a loading control. **(E)** BDQ-CFZ combination reduced OCR compared to the single treatments. OCR was recorded for 36 h with the Resipher™ system after treatments with CFZ and BDQ at the indicated concentrations: BDQ (10 and 15 μM); CFZ (2.5 nM). **(F)** Mitochondrial superoxide generation in AMO-1 cells was measured by flow cytometry 16 and 24 hours post-treatment with CFZ (2.5 nM), BDQ (5 μM), or the combination. Data are shown as mean ± SD of three independent experiments. Statistical significance was determined by one-way ANOVA (****$p < 0.0001$).

**Figure S3. RNA-Sequencing Time Course and Gene Expression Validation Following BDQ-CFZ Treatment. (A)** Schematic of the RNA-sequencing time course. AMO-1 cells were treated with DMSO, CFZ (2.5 nM), BDQ (5 μM), or the combination (BDQ-CFZ) in biological triplicates. Cells were harvested at 6, 12, and 18 hours post-treatment for RNA-seq analysis. Cell viability was concurrently monitored at 6, 12, 18, 36, and 120 hours post-treatment. **(B)** Time-course analysis of cell viability measured by PI staining and flow cytometry. Data represent the mean ± SD of three independent experiments. **(C)** Bar graph summarizing the number of significantly upregulated and downregulated genes (relative to DMSO) at each time point: 133, 637, and 1,087 genes were upregulated, while 124, 682, and 899 were downregulated at 6, 12, and 18 hours, respectively. **(D-E)** Heatmaps showing RT-qPCR validation of selected **(D)** upregulated and **(E)** downregulated genes identified in the RNA-seq dataset after BDQ-CFZ co-treatment. **(F)** Cell cycle analysis of AMO-1 cells 24 hours post-treatment with BDQ (5 μM), CFZ (2.5 nM), or the combination. Cell cycle phases were determined by PI staining and flow cytometry. Data represent the mean of three independent experiments.

**Figure S4. BDQ Does Not Interfere with Trypsin-like and Caspase-like Proteasome Activities and Does Not Synergize with Non-proteasome-Targeting Therapies (A-B)** Caspase-like **(A)** and trypsin-like **(B)** proteasome activities were measured in AMO-1, KMS-11, RPMI-8226, MOLP-8, and U-266 cell lines following 5-hour treatment with BDQ (5 μM), CFZ (2.5 nM), or their combination. **(C-D)** Longitudinal cell viability analysis of Ub-G76V-GFP-expressing AMO-1 and AMO-1$^{BTZ-R}$ cells treated with BDQ (5 μM), CFZ (2.5 nM for AMO-1 and 12.5 nM for AMO-1$^{BTZ-R}$), or the combination. Viability was assessed by TMRM staining and flow cytometry at multiple time points. Results represent the mean ± SD of three independent experiments. **(E)** BDQ does not synergize with non-proteasome–targeting therapies. Heatmap of AMO-1 cell viability following co-treatment with BDQ (5 μM) and escalating doses of Dexamethasone, Venetoclax, Lenalidomide, Melphalan, or Doxorubicin. Cell viability was assessed using CCK-8 assay 72 hours post-treatment.

**Figure S5. BDQ Enhances the Activity of Multiple Proteasome Inhibitors in Both PI-Sensitive and -Resistant MM Cell Lines. (A)** Dose–response curves for AMO-1, AMO-1$^{CFZ-R}$, AMO-1$^{BTZ-R}$, RPMI-8226, RPMI-8226$^{CFZ-R}$, RPMI-8226$^{BTZ-R}$ cell lines treated with increasing concentrations of bortezomib (BTZ), ixazomib (IXA), marizomib (MAR), or carfilzomib (CFZ), with or without 10 μM BDQ. **(B)** Dose–response curves for the same panel of MM cell lines treated with increasing concentrations of TAK-243 in the presence or absence of 10 μM BDQ. Cell viability was assessed 72 hours post-treatment by CellTiter-Glo® Luminescent. **(C)** Summary table reporting the Bliss synergy scores (BS) for each drug combination across the tested cell lines. Green highlights indicate high synergistic interactions (BS > 10), yellow indicates moderate synergistic interactions (1 > BS > 10), while red indicates the absence of synergy (BS < 1).

**Figure S6. Higher Expression of ATP5F1C and ATP5F1B is not Correlated to Worst Prognosis in MM Patients. (A-B)** Analysis of MM patient datasets (CoMMpass IA22) showed that higher expression levels of neither ATP5F1A nor ATP5F1B were associated with poorer overall survival **(A-B)** or progression-free survival **(C-D)**, respectively.

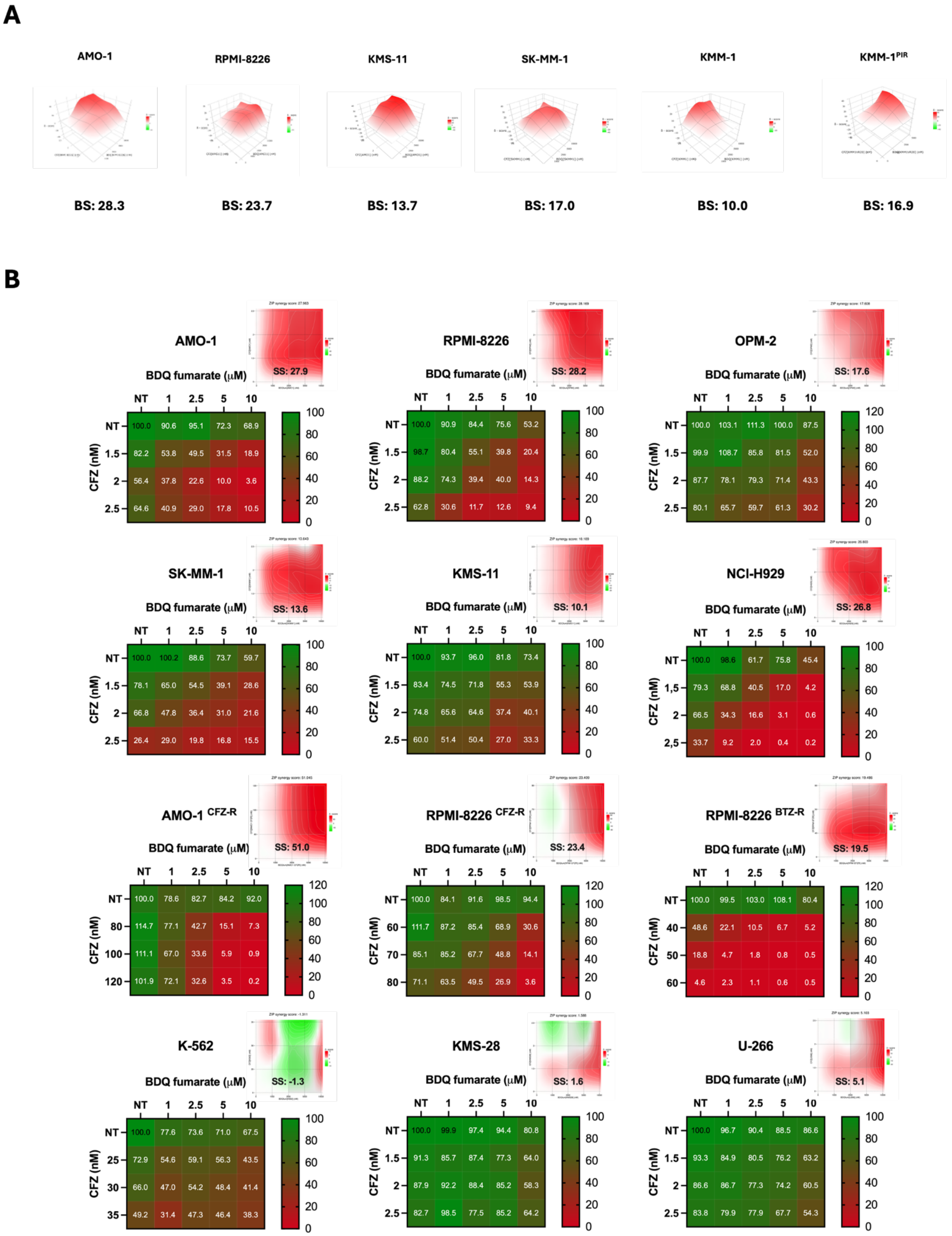
A
AMO-1
RPMI-8226
KMS-11
SK-MM-1
KMM-1
KMM-1PIR
BS: 28.3
BS: 23.7
BS: 13.7
BS: 17.0
BS: 10.0
BS: 16.9
B
AMO-1
SS: 27.9
RPMI-8226
SS: 28.2
OPM-2
SS: 17.6
SK-MM-1
SS: 13.6
KMS-11
SS: 10.1
NCI-H929
SS: 26.8
AMO-1 CFZ-R
SS: 51.0
RPMI-8226 CFZ-R
SS: 23.4
RPMI-8226 BTZ-R
SS: 19.5
K-562
SS: -1.3
KMS-28
SS: 1.6
U-266
SS: 5.1
BDQ fumarate (μM)
CFZ (nM)
NT
1
2.5
5
10

**Figure S1**

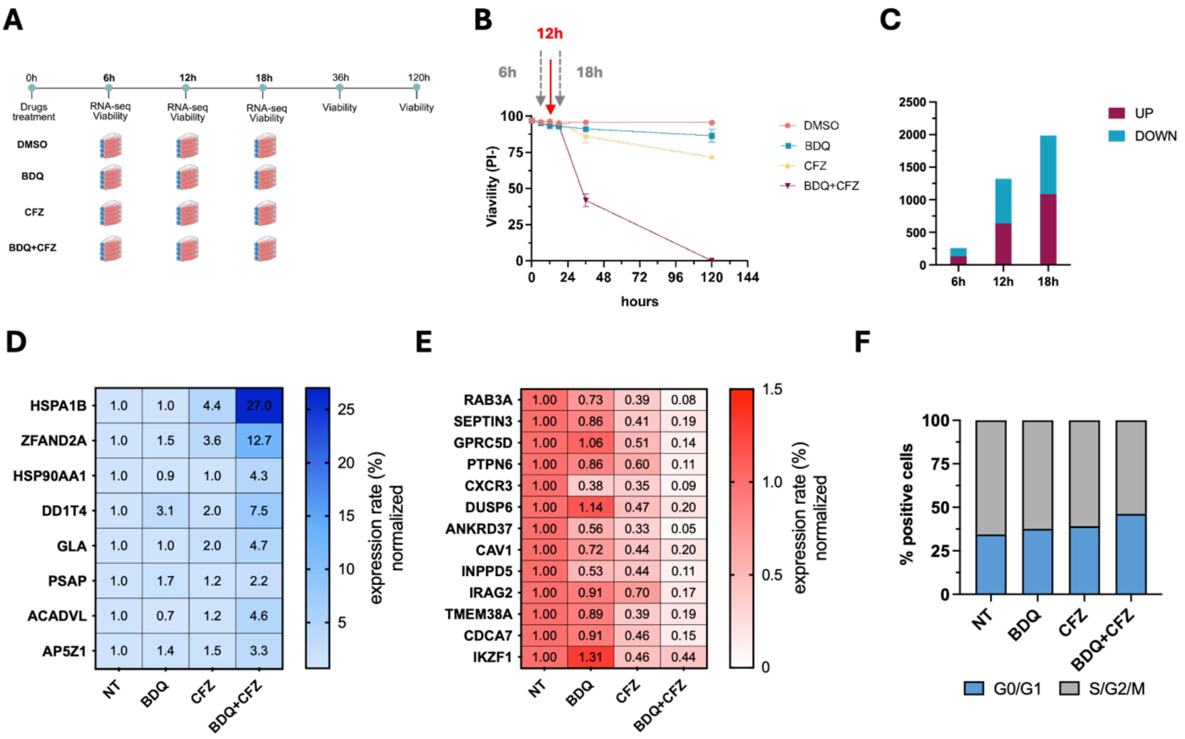
A
0h
6h
12h
18h
36h
120h
Drugs treatment
RNA-seq Viability
RNA-seq Viability
RNA-seq Viability
Viability
Viability
DMSO
BDQ
CFZ
BDQ+CFZ
B
12h
6h
18h
Viavility (PI-)
hours
DMSO
BDQ
CFZ
BDQ+CFZ
C
UP
DOWN
6h
12h
18h
D
HSPA1B 1.0 1.0 4.4 27.0
ZFAND2A 1.0 1.5 3.6 12.7
HSP90AA1 1.0 0.9 1.0 4.3
DD1T4 1.0 3.1 2.0 7.5
GLA 1.0 1.0 2.0 4.7
PSAP 1.0 1.7 1.2 2.2
ACADVL 1.0 0.7 1.2 4.6
AP5Z1 1.0 1.4 1.5 3.3
NT
BDQ
CFZ
BDQ+CFZ
expression rate (%) normalized
E
RAB3A 1.00 0.73 0.39 0.08
SEPTIN3 1.00 0.86 0.41 0.19
GPRC5D 1.00 1.06 0.51 0.14
PTPN6 1.00 0.86 0.60 0.11
CXCR3 1.00 0.38 0.35 0.09
DUSP6 1.00 1.14 0.47 0.20
ANKRD37 1.00 0.56 0.33 0.05
CAV1 1.00 0.72 0.44 0.20
INPPD5 1.00 0.53 0.44 0.11
IRAG2 1.00 0.91 0.70 0.17
TMEM38A 1.00 0.89 0.39 0.19
CDCA7 1.00 0.91 0.46 0.15
IKZF1 1.00 1.31 0.46 0.44
NT
BDQ
CFZ
BDQ+CFZ
expression rate (%) normalized
F
% positive cells
NT
BDQ
CFZ
BDQ+CFZ
G0/G1
S/G2/M

**Figure S3**

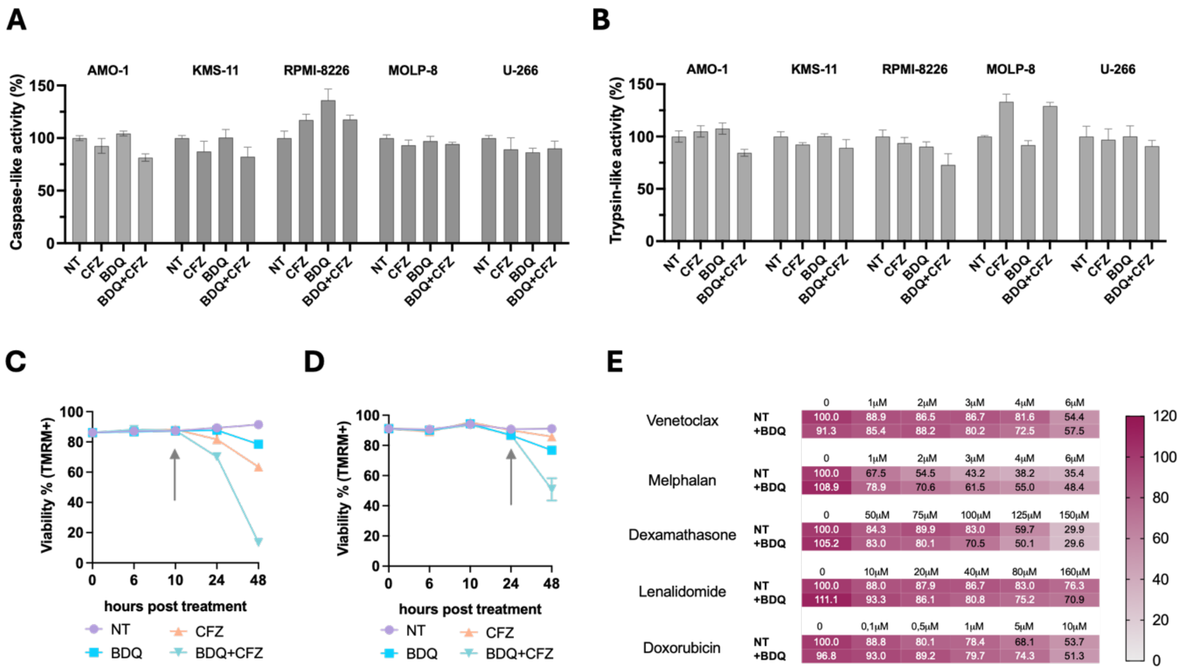
A
AMO-1
KMS-11
RPMI-8226
MOLP-8
U-266
Caspase-like activity (%)
150
100
50
0
NT
CFZ
BDQ
BDQ+CFZ
B
AMO-1
KMS-11
RPMI-8226
MOLP-8
U-266
Trypsin-like activity (%)
C
Viability % (TMRM+)
hours post treatment
0
6
10
24
48
NT
CFZ
BDQ
BDQ+CFZ
D
Viability % (TMRM+)
hours post treatment
NT
CFZ
BDQ
BDQ+CFZ
E
Venetoclax
NT 100.0 88.9 86.5 86.7 81.6 54.4
+BDQ 91.3 85.4 88.2 80.2 72.5 57.5
0 1µM 2µM 3µM 4µM 6µM
Melphalan
NT 100.0 67.5 54.5 43.2 38.2 35.4
+BDQ 108.9 78.9 70.6 61.5 55.0 48.4
0 1µM 2µM 3µM 4µM 6µM
Dexamathasone
NT 100.0 84.3 89.9 83.0 59.7 29.9
+BDQ 105.2 83.0 80.1 70.5 50.1 29.6
0 50µM 75µM 100µM 125µM 150µM
Lenalidomide
NT 100.0 88.0 87.9 86.7 83.0 76.3
+BDQ 111.1 93.3 86.1 80.8 75.2 70.9
0 10µM 20µM 40µM 80µM 160µM
Doxorubicin
NT 100.0 88.8 80.1 78.4 68.1 53.7
+BDQ 96.8 93.0 89.2 79.7 74.3 51.3
0 0,1µM 0,5µM 1µM 5µM 10µM
120
100
80
60
40
20
0

**Figure S4**

A

B

C

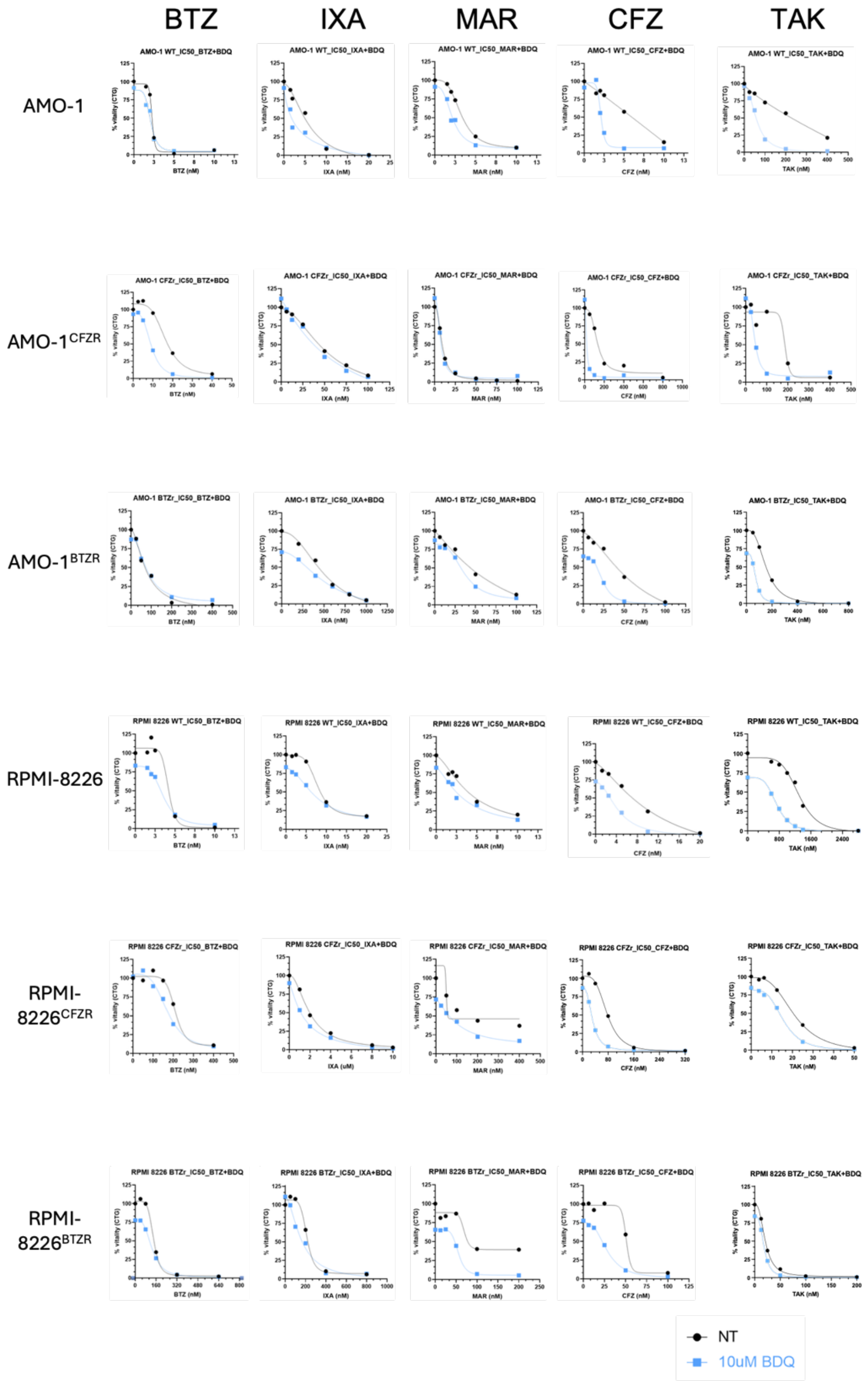


| AMO-1 WT | BS |
|---|---|
| BTZ | 5.5 |
| IXA | 21.1 |
| MAR | 14.4 |
| TAK | 29.5 |
| CFZ | 19.3 |

| AMO-1$^{CFZR}$ | BS |
|---|---|
| BTZ | 21.8 |
| IXA | 10.6 |
| MAR | - 3.5 |
| TAK | 34.3 |
| CFZ | 49.6 |

| AMO-1$^{BTZR}$ | BS |
|---|---|
| BTZ | -7.9 |
| IXA | -1.9 |
| MAR | 2.4 |
| TAK | 13.6 |
| CFZ | 7.0 |

| RPMI WT | BS |
|---|---|
| BTZ | 8.2 |
| IXA | 2.7 |
| MAR | 4.2 |
| TAK | 27.9 |
| CFZ | 9.2 |

| RPMI$^{CFZR}$ | BS |
|---|---|
| BTZ | 13.6 |
| IXA | 7.4 |
| MAR | 10.2 |
| TAK | 7.9 |
| CFZ | 22.6 |

| RPMI$^{BTZR}$ | BS |
|---|---|
| BTZ | 3.3 |
| IXA | 20.5 |
| MAR | 5.8 |
| TAK | 3.6 |
| CFZ | 16.6 |

**Figure S5**

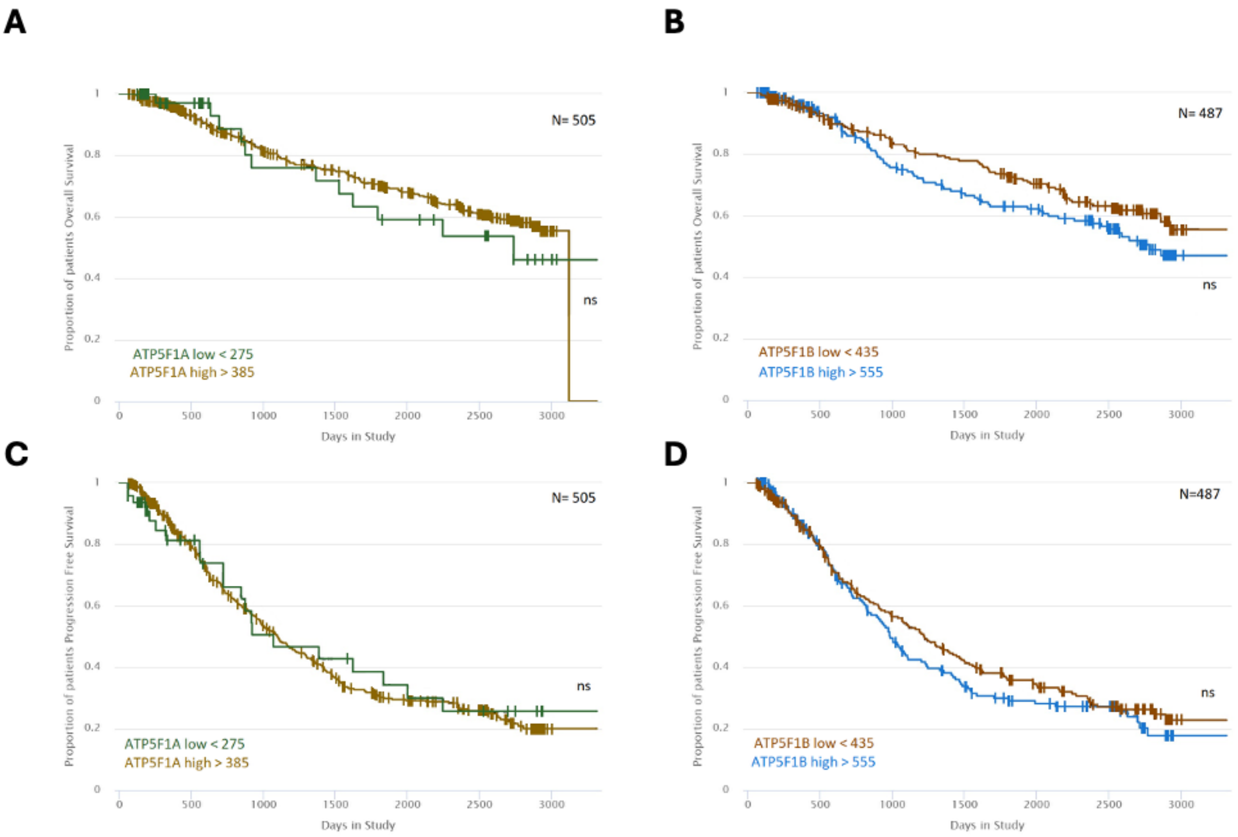
A
B
C
D
N= 505
ns
ATP5F1A low < 275
ATP5F1A high > 385
Proportion of patients Overall Survival
Days in Study
N= 487
ns
ATP5F1B low < 435
ATP5F1B high > 555
Proportion of patients Progression Free Survival
N=487

**Figure S6**

### SUPPLEMENTAL TABLES AND LEGENDS

**Table S1.** Top 14 compounds from the secondary drug screen showing synergy with CFZ in five MM cell lines. The score indicates the number of cell lines (out of five) in which synergy was observed.

| Target | Pathway | Drug | Score |
|---|---|---|---|
| NK1R | GPCR | Aprepitant | 5/5 |
| ATP synthase | Metabolism | Bedaquiline | 5/5 |
| NK1R | GPCR | Rolapitant | 5/5 |
| EHMT2 | Epigenetics | UNC0642 | 5/5 |
| SMYD2 | Epigenetics | LLY-507 | 4/5 |
| BMP-1 | ECM modeling | UK 383367 | 4/5 |
| EGFR | Protein Tyrosine Kinase | AST-1306 | 3/5 |
| NS3 (HCV) | Proteases | Asunaprevir | 3/5 |
| ATP synthase | Metabolism | Bedaquiline fumarate | 3/5 |
| GLS1 | Proteases | BPTES | 3/5 |
| NS5B (HCV) | Proteases | Dasabuvir | 3/5 |
| STAT3 | JAK/STAT | HO-3867 | 3/5 |
| ALK2-ALK3 | TGF-beta/Smad | LDN-193189 2HCl | 3/5 |
| SETD7 | Epigenetics | PFI-2 HCl | 3/5 |

**Table S2.** Cytogenetic and clinical features of MM patients included in ex vivo assays.

| N. | SMM vs NDMM vs RRMM | Date of birth | Del (17p) | T(4;14) | T(14;16) | Gain/Amp (1q) | Del(1p) | T(11;14) | Del(13q) | Disease isotype | ISS at diagnosis | Number of prior lines | Drug exposure | Drug refractoriness |
|---|---|---|---|---|---|---|---|---|---|---|---|---|---|---|
| 8 | NDMM | 29/07/1944 | neg | neg | neg | NV | NV | NV | NV | FLC k | N.A. | 0 | N.A. | N.A. |
| 9 | SMM | 03/09/1959 | neg | neg | neg | NV | NV | NV | NV | IgG l | N.A. | 0 | N.A. | N.A. |
| 22 | NDMM | 08/04/1961 | neg | pos | neg | pos | neg | neg | pos | IgG l | 1 | 0 | N.A. | N.A. |
| 23 | NDMM | 07/05/1962 | neg | neg | neg | neg | neg | neg | pos | IgG k | 3 | 0 | N.A. | N.A. |
| 28 | NDMM | 30/01/1957 | neg | neg | neg | neg | neg | pos | pos | IgG k | 2 | 0 | N.A. | N.A. |
| 30 | NDMM | | pos | neg | neg | pos | neg | NV | NV | IgG k | 1 | 0 | N.A. | N.A. |
| 34 | SMM | 21/03/1950 | neg | neg | neg | pos | neg | neg | pos | IgA k | N.A. | 0 | N.A. | N.A. |
| 36 | SMM | 04/09/1935 | pos | pos | neg | pos | neg | neg | pos | IgG l | N.A. | 0 | N.A. | N.A. |
| 43 | NDMM | 11/04/1947 | neg | pos | NV | NV | NV | NV | pos | IgG l | 3 | 0 | N.A. | N.A. |
| 45 | RRMM | 11/6/1949 | pos | NV | NV | neg | neg | NV | pos | IgG k | 2 | 2 | BTZ, THAL, MEL, CFZ, LENA | BTZ, THAL |
| 52 | RRMM | 23/04/1954 | pos | NV | NV | pos | neg | NV | pos | FLC l | 3 | 1 | DARA, BTZ, THAL, LENA | DARA, LENA |
| 53 | SMM | 07/09/1970 | neg | neg | pos | neg | neg | NV | NV | IgG k | N.A. | 0 | N.A. | N.A. |
| 56 | SMM | 31/01/1945 | neg | neg | neg | pos | neg | neg | pos | IgG k | N.A. | 0 | N.A. | N.A. |

**Table S3.** Concentrations of bedaquiline (BDQ) and carfilzomib (CFZ) used for MM cell line treatments and cell viability assays.

| Cell line | BDQ (μM) | CFZ (nM) | Cell viability detection method |
|---|---|---|---|
| AMO-1 | 5 | 2 | TMRM |
| SK-MM-1 | 5 | 2.5 | TMRM |
| KMM-1 | 5 | 4 | TMRM |
| RPMI-8226 | 5 | 2.5 | TMRM |
| NCI-H929 | 10 | 5 | TMRM |
| OPM-2 | 5 | 2 | TMRM |
| KMS-34 | 10 | 5 | TMRM |
| MOLP-8 | 5 | 2.5 | TMRM |
| JJN-3 | 10 | 50 | TMRM |
| KMS-18 | 5 | 2.5 | TMRM |
| KMS-11 | 5 | 2.5 | TMRM |
| U-266 | 5 | 2.5 | TMRM |
| LP1 | 2.5 | 2.5 | TMRM |
| KMS-26 | 10 | 2.5 | TMRM |
| KMS-28 | 5 | 2 | TMRM |
| K-562 | 10 | 5 | TMRM |
| AMO-1 $^{BTZ-R}$ | 5 | 12.5 | TMRM |
| AMO-1 $^{CFZ-R}$ | 5 | 100 | TMRM |
| RPMI-8226 $^{BTZ-R}$ | 5 | 40 | TMRM |
| RPMI-8226 $^{CFZ-R}$ | 5 | 50 | TMRM |
| KMM-1 $^{PIR}$ | 10 | 7.5 | TMRM |
| U-266 $^{PIR}$ | 5 | 20 | TMRM |
| 5TGM1 | 15 | 25 | PI |
| ALMC-1 | 10 | 1.5 | PI |
| ALMC-2 | 10 | 2 | PI |

**Table S4.** Concentrations of BDQ and CFZ used for B-cell malignancy cell line treatments and cell viability assays.

| Cell line | BDQ (μM) | CFZ (nM) | Cell viability detection method |
|---|---|---|---|
| DOHH2 | 10 | 2.5 | TMRM |
| OCI-LY8 | 10 | 5 | TMRM |
| SU-DHL-6 | 5 | 1.25 | TMRM |
| SU-DHL-7 | 10 | 5 | TMRM |
| Karpas-422 | 10 | 5 | TMRM |
| U-2932 | 7.5 | 1.25 | TMRM |
| RIVA | 10 | 2.5 | TMRM |
| Namalwa | 10 | 2.5 | TMRM |
| BL-41 | 10 | 1.25 | TMRM |
| HS-SULTAN | 5 | 2.5 | TMRM |
| Raji | 10 | 2.5 | TMRM |
| Daudi | 10 | 2.5 | TMRM |
| Granta-519 | 10 | 1.35 | TMRM |
| Mino | 5 | 2.25 | TMRM |

**Table S5.** Concentrations of BDQ and CFZ used for the treatment of bone marrow–derived mononuclear cells (BMMCs).

| Patient (n°) | BDQ (μM) | CFZ (nM) |
|---|---|---|
| 8 | 5 | 2.5 |
| 9 | 10 | 1.25 |
| 22 | 10 | 1.25 |
| 23 | 10 | 1.25 |
| 28 | 10 | 2.5 |
| 30 | 10 | 2.5 |
| 31 | 10 | 2.5 |
| 34 | 10 | 1.25 |
| 36 | 10 | 1.25 |
| 43 | 10 | 1.25 |
| 52 | 5 | 2.5 |
| 53 | 10 | 2.5 |
| 56 | 10 | 0.625 |

**Table S6.** List of oligonucleotide sequences used in this study.

| Gene | Oligo sequence |
|---|---|
| ATP5F1C (shRNA) | Fw: CCGGCGTCAGTCATTGCCCTTGAATCTCGAGATTCAAGGGCAATGACTGACGTTTTTG<br>Rv: AATTCAAAAACGTCAGTCATTGCCCTTGAATCTCGAGATTCAAGGGCAATGACTGACG |
| ACADVL | Fw: GTTGAGCACGGACGGGTATG<br>Rv: GTTGAGCACGGACGGGTATG |
| ANKRD37 | Fw: GTCCACTTAGCCGCAGGAA<br>Rv: CTGCCTTGTGTAGTGGAGCTT |
| AP5Z1 | Fw: GAGAGATGTCCCCCTCTGACA<br>Rv: ACCTCCTCGTTTCTGTCACC |
| ATP5F1C | Fw: GCCCATCTTTTCCCTTAATACCG<br>Rv: TGCAGCACGTCAGCATCAAT |
| CAV1 | Fw: CGCGACCCTAAACACCTCAA<br>Rv: GCCGTCAAAACTGTGTGTCC |
| CDCA7 | Fw: TCCATGGAAACCTCGTCATCC<br>Rv: CCTACAGCCTTCCCGAACTG |
| CXCR3 | Fw: TGCTAAATGACGCCGAGGTT<br>Rv: GGAGGTACAGCACGAGTCAC |
| DDIT4 | Fw: TTTAGCTCCGCCAACTCTCC<br>Rv: TCAATGAGCAGCTGTTCCGA |
| DUSP6 | Fw: CTGGAACGAGAATACGGGCG<br>Rv: ACACCACAGTTCTTGCCCC |
| GAPDH | Fw: TCTTTTGCGTCGCCAGCCGAG<br>Rv: TGACCAGGCGCCCAATACGAC |
| GLA | Fw: CTCATGGGATTCGCCAGCTA<br>Rv: CAAAACTCCCAGGGAAGCCT |
| GPRC5D | Fw: GTACTCCCATTCAGCCGCAG<br>Rv: TCTTGCTGGGGGCTTAGTTT |
| HSPA1B | Fw: TCAGGCCCTACCATTGAGGA<br>Rv: GAGTCCCAACAGTCCACCTC |
| HSP90AA1 | Fw: GTCCTGTGCGGTCACTTAGC<br>Rv: ACTGGGCAATTTCTGCCTGA |
| HUPO | Fw: GCTTCCTGGAGGGTGTCC<br>Rv: GGACTCGTTTGTACCCGTTG |
| IKZF1 | Fw: TGGCAGGGCAGAGGGAG<br>Rv: GCATCCATGGTCCTCAGGTT |
| INPPD5 | Fw: CGCCCATATCACCCAAGAAGT<br>Rv: CTGCGTTCTTCCCCAGGTC |
| IRAG2 | Fw: AACACTTGCCAGGCACTTCA<br>Rv: AAGCCATGGTGTTCCCCTTC |
| PSAP | Fw: GTGGACTCCTACCTCCCTGT<br>Rv: GTTGAGAGCAGAGCACACCT |
| PTPN6 | Fw: AACAGCCGTGTCATCGTCAT<br>Rv: AGTAGGGCCCATAAGCACG |
| RAB3A | Fw: AAAAAGTCACCGCCGCTAGG<br>Rv: GATCCGAGGACTCCTTCTGC |
| SEPTIN3 | Fw: CGACACCATCATCGAGCAGA<br>Rv: ATTTGCCCAGTCCACTCTGG |
| TMEM38A | Fw: ATTGCAACTGGGTGGGTCAA<br>Rv: TGTGCAGGATCTCGTTGGTC |
| ZFAND2A | Fw: ATACCAGACGTGGTGGTTGG<br>Rv: CAGCCCTCTTTTGAGCAACG |

**Table S7.** Primary antibodies employed in this study.

| Antibody | Species | Source |
|---|---|---|
| 53BP1 | rabbit | #4937, Cell Signaling Technology |
| p-ATM | rabbit | #5883 Cell Signaling Technology |
| ATM | rabbit | #28735 Cell Signaling Technology |
| α-Tubulin | mouse | clone B-5-1-2, T5168, Sigma-Aldrich |
| β-Actin | mouse | #sc69879, SCB |
| ATP5F1C | rabbit | AB119686-1001 Abcam |
| p-H2A.X (Ser139) | rabbit | #9718T Cell Signaling Technology |
| RPA32/RPA2 | rabbit | #52448 Cell Signaling Technology |
| p-RPA32/RPA2 (Ser8) | rabbit | #83745 Cell Signaling Technology |
| Cleaved PARP | rabbit | #9532 Cell Signaling Technology |
| PARP | rabbit | #9542 Cell Signaling Technology |
| Cleaved Caspase 3 | rabbit | #9664T Cell Signaling Technology |
| Caspase 3 | rabbit | #14220T Cell Signaling Technology |
| V5 tag | mouse | #sc81594, SBB |
| Vinculin | mouse | #SAB4200080, Sigma-Aldrich |
